# Task-induced 1/f slope modulation as a paradigm-independent marker of cognitive control in multiple sclerosis

**DOI:** 10.1101/2025.04.11.648423

**Authors:** Fahimeh Akbarian, Máté Gyurkovics, Marie B D’hooghe, Miguel D’haeseleer, Guy Nagels, Jeroen Van Schependom

**Affiliations:** Department of Electronics and Informatics (ETRO), Vrije Universiteit Brussel, Brussels, Belgium; AIMS lab, Vrije Universiteit Brussel, Center for Neurosciences, Brussels, Belgium; School of Psychology, University of East Anglia, UK; National MS Center Melsbroek, Melsbroek, Belgium; Center for Neurosciences, Vrije Universiteit Brussel, Brussels, Belgium; UZ Brussel, Department of Neurology, Brussels, Belgium; St Edmund Hall, University of Oxford, Oxford, UK

**Keywords:** Aperiodic 1/f slope, multiple sclerosis, magnetoencephalography (MEG), excitation/inhibition (E/I) balance, aperiodic activity, auditory oddball paradigm

## Abstract

Multiple sclerosis (MS) is a chronic neuro-degenerative and inflammatory disease causing motor, sensory, and cognitive deficits, including impairments in working memory and attention. These cognitive deficits may arise from an imbalance between excitatory and inhibitory neural activity due to synaptic loss. Recent studies suggest that the aperiodic 1/f slope, a neural marker reflecting excitation/inhibition (E/I) balance, could serve as a biomarker for cognitive control. This study examines 1/f slope modulation during cognitive tasks in people with MS and healthy controls to investigate its potential as a paradigm-independent marker of cognitive control.

We analyzed the Magnetoencephalography (MEG) data collected from 126 participants: 44 healthy controls (HCs), 61 people with MS not treated with benzodiazepines (pwMS(BZDn)), and 21 pwMS treated with benzodiazepines (pwMS(BZDp)). Participants performed an auditory oddball task and a visual-verbal n-back working memory task. After preprocessing MEG data, we used the FOOOF algorithm to extract the aperiodic 1/f slope from power spectral densities across 42 cortical parcels.

Through this analysis, we observed significant increases in the 1/f slope following stimulus onset for all stimulus types, with more pronounced modulation for non-standard stimuli (targets and distractors), especially within the temporal cortex. Group comparisons revealed less slope modulation in pwMS(BZDp) compared to HCs during distractor stimuli, indicating impaired inhibitory control linked to benzodiazepine treatment. Positive correlations of 1/f slope modulation across auditory oddball and n-back tasks were observed in HCs and pwMS(BZDn), highlighting a consistent, paradigm-independent mechanism.

Taken together, these findings demonstrate that the aperiodic 1/f slope is a sensitive, paradigm-independent marker of cognitive control and E/I balance. Reduced modulation in response to distractors among pwMS(BZDp) highlights benzodiazepine-related disruptions in inhibitory neural processes underlying cognitive deficits. These findings underscore the value of aperiodic spectral measures to deepen understanding and potentially guide therapeutic interventions targeting cognitive impairments in MS.

**Highlights:**

- The 1/f slope was significantly modulated during an auditory oddball task across all stimulus types in healthy controls and people with multiple sclerosis (pwMS).
- This modulation was more pronounced for non-standard stimuli (distractor and target) than for standard stimuli.
- Healthy controls showed a larger 1/f slope modulation (suggesting more inhibition) following distractor trials compared to target trials while, this pattern was reversed in pwMS.
- This modulation significantly correlated with visuospatial memory, as measured by the BVMT-R, in pwMS not treated with benzodiazepines.
- We further correlated the 1/f slope modulation during the auditory oddball task with the 1/f slope modulation previously described during a working memory task and observed strong correlations (r=0.4-0.6, p<0.001) across paradigms.

## 1. Introduction

Multiple sclerosis (MS) is a chronic autoimmune disorder characterized by widespread demyelination and neurodegeneration within the central nervous system. Beyond motor and sensory deficits, MS is frequently associated with significant cognitive impairments, particularly in working memory (WM), attention, information processing speed, and executive functions (Benedict et al., 2004; Rao et al., 1991). Despite extensive research, the neural mechanisms underlying cognitive impairments in MS remain poorly understood.

One characteristic of MS pathology is synaptic loss, particularly affecting inhibitory synapses, leading to disruption of the balance between excitation and inhibition (E/I) in neural circuits (Huiskamp et al., 2022; Zoupi et al., 2021). This imbalance leads to reduced network stability and impaired cognitive control, which may underline the observed cognitive deficits in people with MS (pwMS) (Huiskamp et al., 2022). One possible approach to understanding the neural substrates of cognitive impairments in MS is to investigate the imbalance in the dynamic interaction between excitatory and inhibitory neuronal circuits.

Traditional neurophysiological studies aimed at understanding cognitive impairment in MS have long focused on oscillatory neural activity by analyzing rhythms within specific frequency bands (Keune et al., 2017; Khan et al., 2021; Simon et al., 2023). While these oscillatory measures provide valuable insights, they only capture one aspect of the rich dynamics of neural activity. In recent years, there has been growing interest in the non-oscillatory or aperiodic component of the neural power spectrum, often modelled as 1/f activity, as a complementary approach to understanding neural function. The 1/f activity captures the broadband shape of the power spectrum where power typically decreases as a function of frequency (or f) according to a power-law distribution, typically approximated by 1/f^x^ (Donoghue et al., 2020). Unlike narrowband oscillatory measures, the aperiodic 1/f component reflects scale-free properties across a wide frequency range, offering a window into the global state of neural excitability and the balance between excitation and inhibition (Gao et al., 2017; He et al., 2010). Recent theoretical and methodological advances have further bolstered the evidence supporting its functional relevance in elucidating brain dynamics and human behavior (Donoghue et al., 2020; Gyurkovics et al., 2022; Kałamała et al., 2023; Voytek et al., 2015; Waschke et al., 2021).

Central to this approach is the 1/f slope, or exponent, which quantifies the rate at which power declines with increasing frequency within the power spectral density of neural data. Evidence suggests that the 1/f slope serves as an index of E/I balance in the brain (Gao et al., 2017). A steeper slope indicates a higher level of inhibition, while a flatter slope indicates increased excitation (Akbarian et al., 2023; Chini et al., 2022; Gao et al., 2017). Importantly, this metric is not merely a reflection of noise in neural activity but is associated with cognitive functioning. Task-induced changes in the 1/f slope have been linked to cognitive demands such as attention and working memory load, highlighting its role as an index of cognitive control (Akbarian et al., 2024; Gyurkovics et al., 2022; Lu et al., 2024; Waschke et al., 2021). Therefore, understanding how the 1/f slope adapts to cognitive demand provides valuable insights into the neural substrates of cognition and their disruption in MS.

In this study, we first explore event-related 1/f slope modulation during an auditory oddball task in pwMS and healthy controls (HCs). The auditory oddball task probes selective attention by requiring participants to discriminate between standard and deviant (target/distractor) stimuli (Squires et al., 1975). Based on previous findings linking steeper spectral slopes to increased cognitive demands (Gyurkovics et al., 2022; Lu et al., 2024), we hypothesize that the 1/f slope modulation (post-event steepening) will be more pronounced for non-standard (target/distractor) stimuli than standard stimuli. We also hypothesize that pwMS will exhibit lower task-related 1/f slope modulation than HCs during distractor trials, as we previously observed during a working memory task, potentially due to a disruption in their cortical E/I balance (Akbarian et al., 2024).

We further aim to investigate whether the 1/f slope modulation remains consistent across different tasks. Specifically, we examine whether the modulation of the 1/f slope observed during the auditory oddball task correlates with that observed in a visual-verbal n-back task, as suggested by our previous findings (Akbarian et al., 2024). The n-back task, a widely used measure of working memory, involves monitoring and updating information with varying levels of difficulty (0-back, 1-back, 2-back) (Costers et al., 2020). By assessing 1/f slope modulation in both tasks, we aim to determine whether these changes are task-specific or indicative of a shared, paradigm-independent cognitive control mechanism. In light of recent findings showing that aperiodic modulations are sensitive to multiple types of cognitive demand (working memory, task switching, and inhibitory control;(Lu et al., 2024)), we predicted results to be in favor of the latter alternative. Furthermore, demonstrating correlations across the two tasks would support the notion that the 1/f slope reflects a shared neurophysiological mechanism potentially a trait-like indicator of top-down control capacity rather than being solely task-specific.

## 2. Methods

### 2.1. Participants

MEG data were collected from 126 participants during an auditory oddball task, including 21 people with MS (pwMS) receiving benzodiazepine treatment (pwMS(BZDp)), 61 pwMS not receiving benzodiazepines (pwMS(BZDn)), and 44 healthy controls (HCs). Additionally, 101 of these participants (34 HCs, 17 pwMS(BZDp), 50 pwMS(BZDn)) also completed a visual-verbal n-back task. Given our recent findings (Akbarian et al., 2023) that benzodiazepines influence the 1/f spectral slope, we categorized the pwMS group based on their benzodiazepine treatment status. The participants were between 18 and 65 years old (mean = 47.85, SD = 10.64). All pwMS were recruited from the National MS Center Melsbroek and were diagnosed with multiple sclerosis according to the revised McDonald’s criteria (Polman et al., 2011). They had an Expanded Disability Status Scale (EDSS) score of 6 or lower (Kurtzke, 1983) to ensure the feasibility of patient participation in data acquisition. Exclusion criteria included a history of relapses or corticosteroid treatment within the previous six weeks, as well as the presence of a pacemaker, dental wires, major psychiatric disorders, or epilepsy. All participants provided written informed consent before participation. The study received ethical approval from the local ethics committees of the University Hospital Brussels (Commissie Medische Ethiek UZ Brussel, B.U.N. 143201423263, 2015/11) and the National MS Center Melsbroek (2015-02-12).

### 2.2. Data acquisition

The MEG data was collected at the CUB Hôpital Erasme (Brussels, Belgium) on an Elekta Neuromag Vectorview scanner (Elekta Oy, Helsinki, Finland) for the first 30 pwMS and 14 healthy subjects and the rest of subjects were scanned using an upgraded scanner, Elekta Neuroimage Triux scanner (MEGIN, Croton Healthcare, Helsinki, Finland). Both MEG scanners used a sensor layout with 102 triple sensors, each consisting of one magnetometer and two orthogonal planar gradiometers and were placed in a lightweight magnetically shielded room (MaxshieldTM, Elekta Oy, Helsinki, Finland). MEG data were recorded during the auditory oddball and n-back task data. Structural MRI scans were obtained using a 3T Philips MRI system with a T1-weighted sequence at of 1 × 1 × 1 mm³ resolution. The acquisition parameters included a repetition time (TR) of 4.93 ms, a flip angle (FA) of 8°, a field of view (FOV) of 230 × 230 mm², and 310 sagittal slices, achieving a voxel resolution of 0.53 × 0.53 × 0.53 mm³.

### 2.3. MEG processing and parcellation

MEG signals were recorded with a 0.1–330 Hz passband filter at 1 kHz sampling rate. The preprocessing of MEG data began with applying the temporal extension of the signal space separation algorithm (MaxFilter™, Elekta Oy, Helsinki, Finland, version 2.2, default parameters) to eliminate external interferences and correct for head movements (Taulu et al., 2005). Subsequent preprocessing was conducted using the Oxford Software Library (OSL) pipeline, which integrates tools from FSL, SPM12 (Wellcome Trust Centre for Neuroimaging, University College London), and FieldTrip (Oostenveld et al., 2011).

First, we applied a finite impulse response (FIR) anti-aliasing low-pass filter with a cut-off frequency at 125 Hz, then the data were downsampled to 250 Hz. Coregistration with the subject’s T1-weighted MRI was performed automatically using OSL’s RHINO algorithm (https://github.com/OHBA-analysis). Head shape points were aligned with the scalp surface extracted via FSL’s BETSURF and FLIRT (Jenikson, Mark, 2005; Smith, 2002) and subsequently transformed into the common MNI152 space (Mazziotta et al., 1995). The data were then band-pass filtered between 0.1 and 70 Hz, and a 5th-order Butterworth notch filter (49.5– 50.5 Hz) was applied to remove power line noise.

To address artefacts, a semi-automated independent component analysis (ICA) was performed to visually identify and remove ocular and cardiac artefacts, based on the correlation of component time series with electrooculogram (EOG) and electrocardiogram (ECG) signals, respectively. For source reconstruction, a linearly constrained minimum variance (LCMV) beamformer was employed to project MEG data into source space (Oostenveld et al., 2011; Quinn et al., 2018; Woolrich et al., 2011). The source-reconstructed signals were then parcelled using a predefined atlas consisting of 42 cortical parcels (Vidaurre et al., 2018). Within each parcel, the first principal component (PC) of the time series was extracted and used as the representative signal. The parcellation atlas was designed to cover the entire cortex, excluding subcortical regions (Van Schependom et al., 2019; Vidaurre et al., 2018).

### 2.4. Auditory oddball task

The subjects were presented with a series of 400 auditory tones over a total test duration of 8 minutes and 20 seconds. Eighty percent of the tones were categorized as frequent tones (1000 Hz), 10% as target tones (1500 Hz), and 10% as distractor tones (500 Hz). Subjects were instructed to press a button upon detecting a high-pitched target tone amidst the series of standard tones. The interstimulus interval was randomized between 1 and 1.5 seconds. Accuracy was defined as the proportion of correct responses (either correct detections or correct rejections) relative to the total number of trials.

### 2.5. Visual-verbal n-back task

During the MEG acquisition, all participants were asked to perform an n-back task (Costers et al., 2020) with three conditions or levels of working memory load (0, 1 and 2-back). The n-back task includes trials that require a response and those that do not, referred to as “target” trials and “distractor” trials, respectively. The examiner instructed participants to press a button with their right hand when the letter displayed on the screen was the letter X (0-back condition), the same letter as the one before (1-back condition), or the same letter as two letters before (2-back condition). There were 240 stimuli, with 25, 23, and 28 target trials per condition (0-back, 1-back, 2-back), respectively.

### 2.6. Power spectral analysis and estimation of aperiodic components

We calculated the power spectral density (PSD) for each trial and parcel using SciPy’s default Welch function, following the method described in (Akbarian et al., 2024) to extract the 1/f slope modulation. Next, we applied the “Fitting Oscillations and One-Over F” (FOOOF) algorithm (Donoghue et al., 2020) to estimate the 1/f exponent that quantifies the steepness of the 1/f slope. This algorithm iteratively fits Gaussian functions to the periodic components of the PSD and subtracts them, thereby isolating the aperiodic 1/f component. The fitting was performed over 4 Hz to 44 Hz frequency range. In the auditory oddball paradigm, each trial provided 1 second of data (0.5 seconds before and 0.5 seconds after the stimulus), while in the n-back paradigm, we used 1-second windows both before and after the stimulus onset. First, we excluded trials where the subject had pressed the button on the immediately preceding trial, to eliminate any influence of button presses on the pre-stimulus time window. Then power spectra densities were computed separately for 42 brain parcels for each subject, and the corresponding 1/f exponents (or slopes) were extracted using the FOOOF algorithm.

To adjust for event-related fields (ERFs) during the post-stimulus period, we implemented the method described by Gyurkovics et al (Gyurkovics et al., 2022). First, we measured the power spectra from averaged time-domain responses (i.e., the ERFs) for each stimulus type, subject, brain parcel, and time window. These ERF spectra were then subtracted from the total power spectra (i.e., the average of single-trial spectra), effectively isolating induced activity by removing the ERF contribution. We applied the same procedure to correct for ERF-related effects in the power spectra from the n-back task data.

### 2.7. Neuropsychological assessment

Neuropsychological tests were performed on the same day as the MEG recording for all participants. The test battery included the Symbol Digit Modalities Test (SDMT, (A Smith, 1968)) to evaluate information processing speed, the Dutch version of the California Verbal Learning Test (CVLT-II, (Strober et al., 2009)), the Dutch version: VGLT to assess verbal memory. Verbal fluency was measured by the Controlled Oral Word Association Test (COWAT, (Ruff et al., 1996)), and spatial memory was evaluated using the Brief Visuospatial Memory Test (Revised; BVMT-R, (Benedict et al., 1996)). Additionally, Fatigue was assessed by the Fatigue Scale for Motor and Cognitive Function (FSMC, (Penner et al., 2009)) and depression by Beck’s Depression Inventory (BDI, (Beck, 1961)).

### 2.8. Statistics

All statistical analyses were conducted using non-parametric methods, which do not assume a specific data distribution. Paired comparisons were assessed using the Wilcoxon signed-rank test, while non-paired comparisons were analyzed using the Mann-Whitney U test. To account for multiple comparisons, Benjamini-Yekutieli false discovery rate (FDR) correction (Benjamini & Hochberg, 1995) was applied to adjust p-values, with the false positive rate set to 5%. Effect sizes for both within-group and between-group statistical tests are reported as Cohen’s d-values (Cohen, 1988), which are interpreted as follows: 0.1 to 0.3 indicates a small effect, 0.3 to 0.5 is an intermediate effect, and 0.5 or greater is a strong effect. A significant threshold of p < 0.05 was used for all statistical tests.

## 3. Results

### 3.1. Behavioral data

Figure 2 shows the distribution of reaction time and accuracy for different groups. The group comparisons revealed a significant difference in accuracy between healthy controls (HCs) and pwMS(BZDp) patients, with HCs showing better performance (p = 0.007). No significant differences were observed in either accuracy or reaction times for any other between-group comparisons.

**Figure 1.**
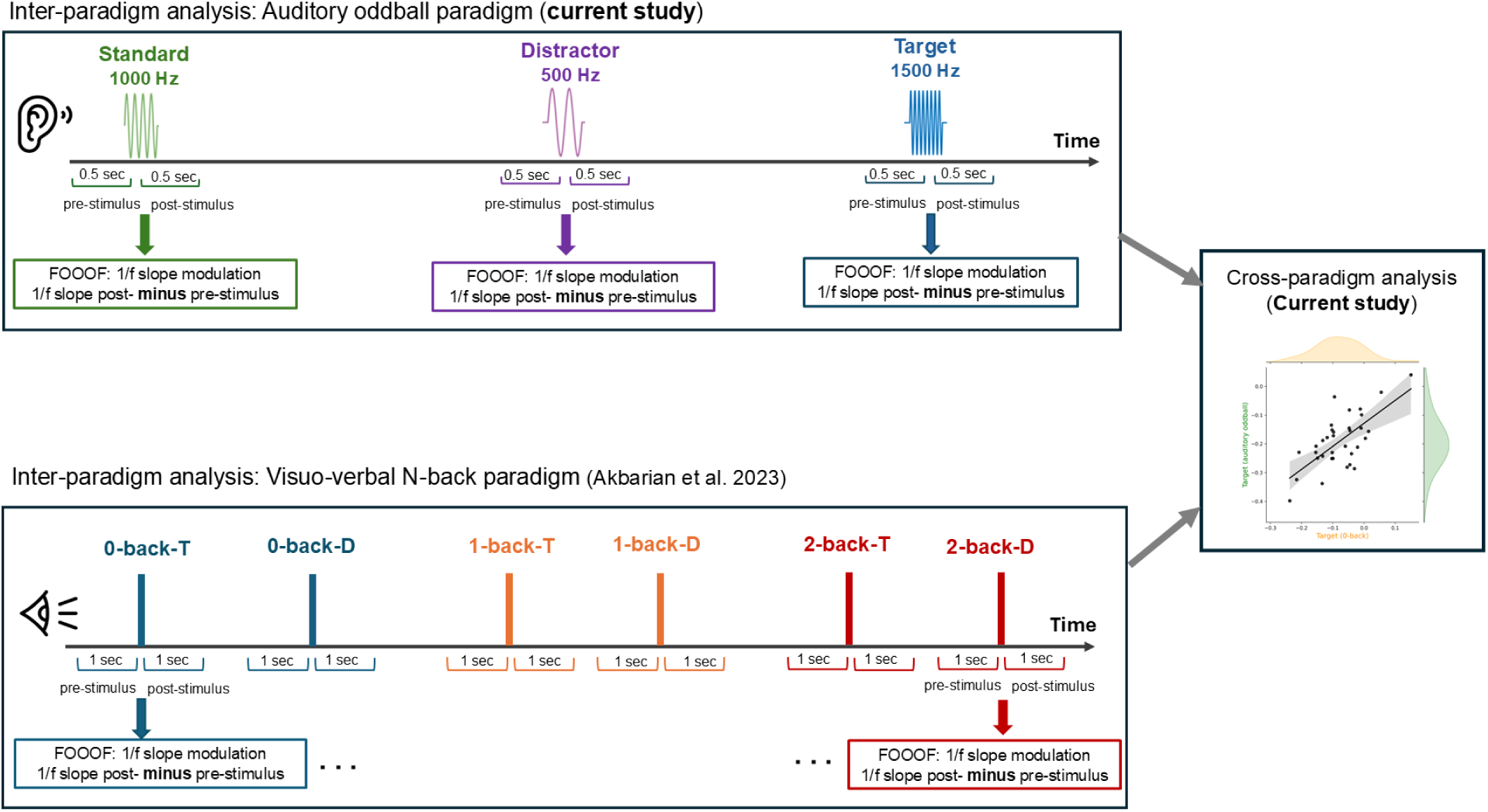
Study pipeline. The task-induced modulation of the 1/f slope during a working memory task was previously described (Akbarian et al., 2024). In the current study, we analyzed the task-induced modulation of the 1/f spectral slope during an auditory oddball paradigm. Findings from both analyses were used to investigate cross-paradigm modulation of the 1/f slope.

**Figure 2.**
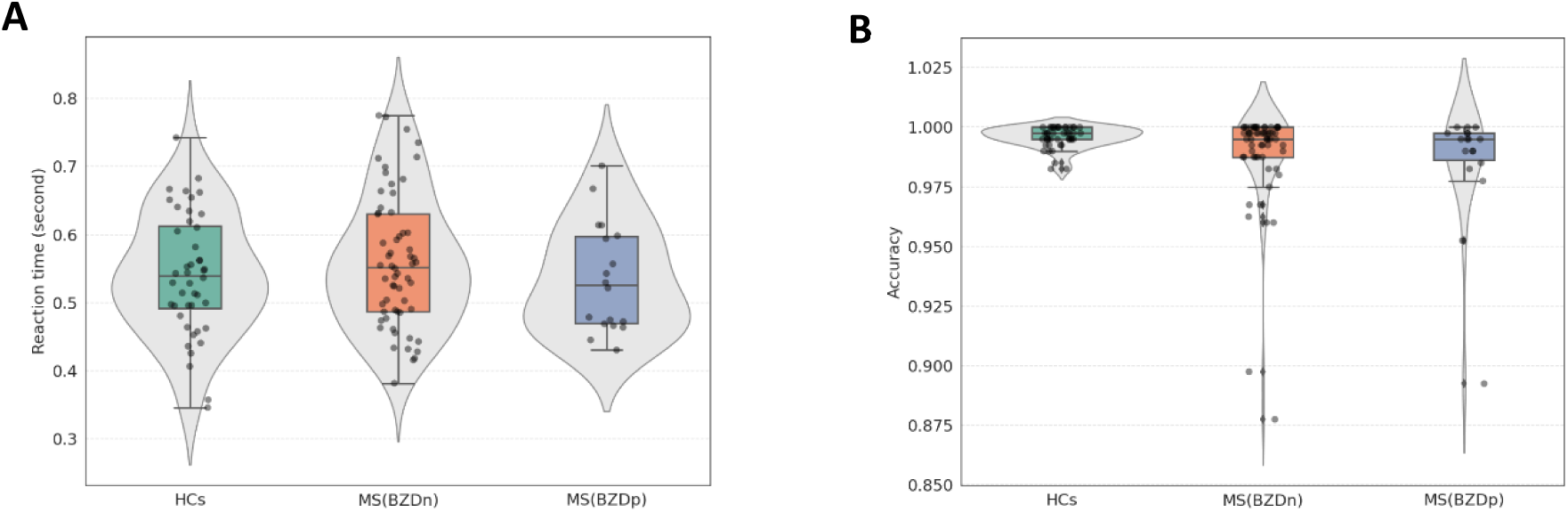
The distribution of the **A)** reaction times and **B)** accuracy for three groups (HCs, pwMS(BZDn), pwMS(BZDp)) are shown respectively. The Mann-Whitney tests were used to compare the reaction times and accuracy between different groups.

### 3.2. The task-induced 1/f slope modulation above and beyond the ERFs

Across all groups, we observed a robust and statistically significant increase in the 1/f slope following stimulus onset for each stimulus type (standard, target, and distractor) compared to the pre-stimulus baseline. The Wilcoxon Signed-Rank Test results confirmed this effect within each group. As shown in Figure 3 for healthy controls, the increase was significant for standard (W = 189, Z= -3.5, p < 0.0001, d = 0.53), distractor (W = 12, Z = -5.6, p < 0.0001, d = 0.84) and target stimuli (W = 18, Z = -5.56, p < 0.0001, d = 0.83). Similarly, in pwMS (BZDn), significant effects were observed for standard (W = 542, Z = -2.89, p = 0.003, d = 0.37), distractor (W = 91, Z = -6.13, p < 0.0001, d = 0.78) and target stimuli (W = 70, Z = -6.28 p < 0.0001, d = 0.80), as it is shown in Figure 4. In pwMS (BZDp), the increase was also significant for standard (W = 55, Z = -2.10, p = 0.03, d = 0.36), distractor (W = 38, Z = -2.69, p = 0.005, d = 0.58) and target stimuli (W = 4, Z= -3.87, p < 0.0001, d = 0.84), see Figure S1.

**Figure 3.**
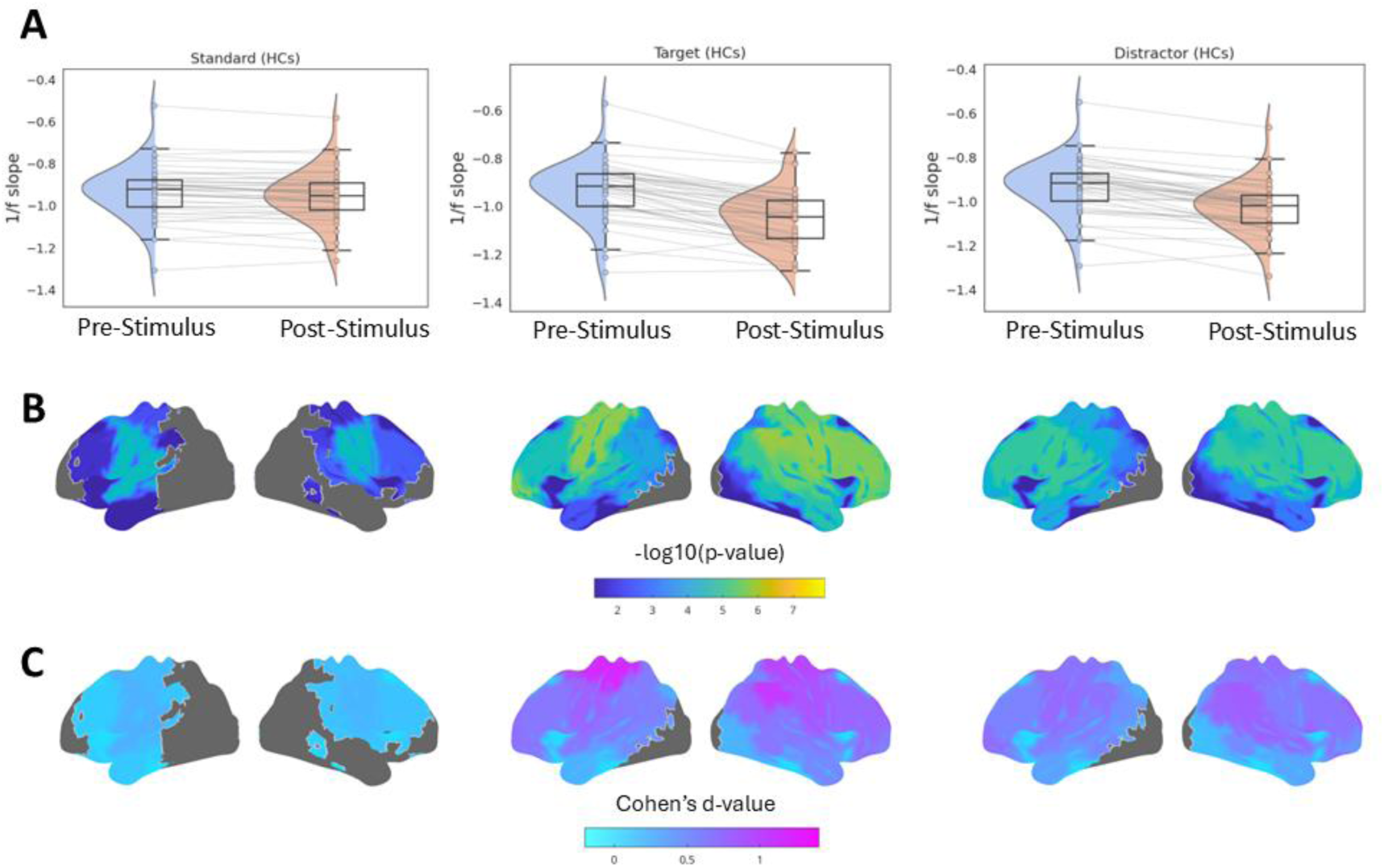
**(A)** Distribution of whole-brain averaged 1/f slope values measured pre- and post-stimulus for all trials in **healthy controls** (HCs). The Wilcoxon signed-rank test yielded p-values < 0.0001 for all within-group comparisons. **(B)** Spatial distribution maps at the parcel level (42 brain parcels) comparing the 1/f slope pre- and post-stimulus, with corrections for multiple comparisons using the FDR method. **(C)** Spatial distribution of effect sizes, represented as Cohen’s d for each parcel-wise comparison.

**Figure 4.**
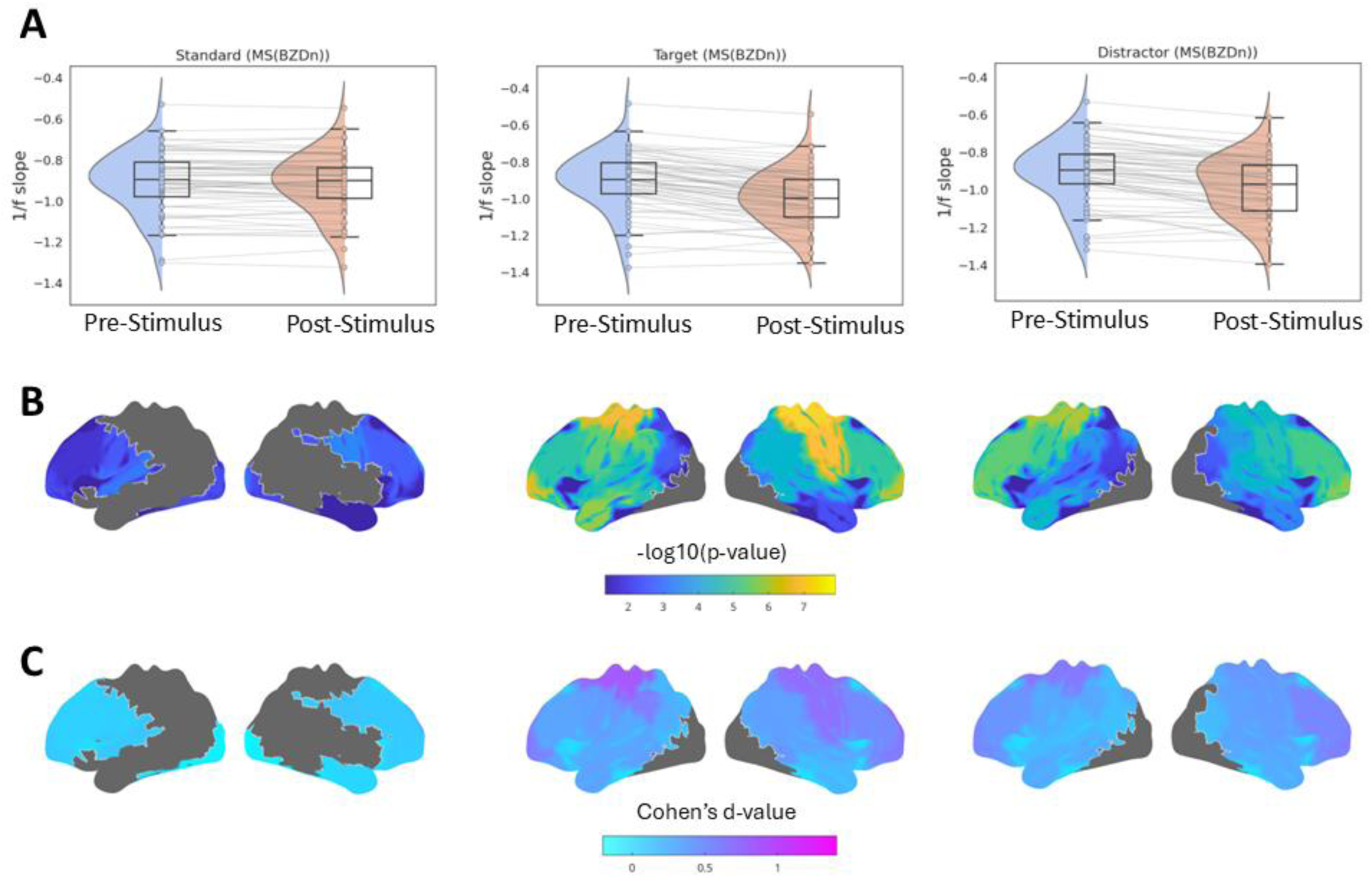
**(A)** Distribution of whole-brain averaged 1/f slope values measured pre- and post-stimulus for all trials in **pwMS(BZDn)**. The Wilcoxon signed-rank test yielded p-values < 0.0001 for all within-group comparisons. **(B)** Spatial distribution maps at the parcel level (42 brain parcels) comparing the 1/f slope pre- and post-stimulus, with corrections for multiple comparisons using the FDR method. **(C)** Spatial distribution of effect sizes, represented as Cohen’s d for each parcel-wise comparison.

Parcel-wise analysis revealed that this effect was most pronounced in the temporal cortex. Notably, the effect was more spatially widespread for non-standard stimuli (target and distractor) than for standard stimuli, indicating greater cortical engagement when participants responded to deviant sounds.

### 3.3. The effect of group and trials on 1/f slope modulation

To further evaluate the effect of group and trial on the amount of modulation in 1/f slope following the stimulus presence, we conducted a two-way ANOVA on the rank-transformed 1/f slope modulation (post-stimulus minus pre-stimulus 1/f slope). The ANOVA was conducted with a group (HCs, pwMS(BZDn), pwMS(BZDp)) as the between-subject factor and trial type (standard, target, distractor) as the within-subject factor. The results of the ANOVA analysis are provided in Table 2.

**Table 1.** Description of subjects. We report the mean values and standard deviations for different clinical parameters. For EDSS, the median and interquartile range (IQR) are shown. The comparisons were performed using permutation testing with N = 5000 for all parameters except for sex, for which a chi-squared test was used. HCs: healthy controls, pwMS(BZDp) and pwMS(BZDn): pwMS with and without benzodiazepines. The “p” in the term for patients treated with benzodiazepines refers to the treatment status, which is positive”. Conversely, “n” in the term for patients not treated with benzodiazepines refers to the treatment status, which is “negative”.

|  | HCs | pwMS sub-groups |  | Group differences (p-values) |  |  |
| --- | --- | --- | --- | --- | --- | --- |
|  | HCs | pwMS(BZDn) | pwMS(BZDp) | HCs vs pwMS(BZDn) | HCs vs pwMS(BZDp) | pwMS(BZDn) vs<br>pwMS(BZDp) |
| *N* | 44 | 61 | 21 | - | - | - |
| *Sex (M/F)* | 18/26 | 20/41 | 2/19 | 0.51 | **0.02** | 0.07 |
| *Age (yrs)* | 47±12 | 48±10 | 47±7 | 0.55 | 0.92 | 0.69 |
| *Education (yrs)* | 15±2 | 14±2 | 13±2 | 0.05 | **0.007** | 0.30 |
| *Disease duration(yrs)* | - | 17±10 | 14±6 | - | - | 0.19 |
| *EDSS median [IQR]* | - | 2.5 [2-4] | 3[2.5-4] | - | - | 0.12 |
| *Cognitive assessment* |  |  |  |  |  |  |
| *SDMT* | 53 ±7 | 49±12 | 46±7 | 0.06 | **0.004** | 0.34 |
| *VGLT* | 65 ±7 | 64±10 | 63±10 | 0.43 | 0.31 | 0.74 |
| *COWAT* | 14±4 | 13±4 | 14±4 | **0.02** | 0.87 | 0.15 |
| *BVMT-R* | 28±5 | 25±7 | 25 ±6 | **0.02** | 0.09 | 0.89 |

**Table 2.** Summary of ANOVA results on the rank-transformed 1/f slope modulation. The effect size is reported as Eta-squared (η²) with a range of 0 (no effect) to > 0.2 (large effect).

|  | *df* | *F* | *p* | *Eta-squared ($\eta^2$)* |
| --- | --- | --- | --- | --- |
| *Groups*<br>[HCs, pwMS(BZDn), pwMS(BZDp)] | 2 | 6.82 | $P < 0.001$ | 0.03 |
| *Trials*<br>[standard, target, distractor] | 2 | 94.31 | $p < 0.0001$ | 0.49 |
| *Groups X trials* | 4 | 0.64 | 0.65 | 0.007 |

A main effect of group was observed, indicating that the three groups (HCs, pwMS(BZDn), pwMS(BZDp)) differed in their overall 1/f slope modulation, F(2, 369) = 6.82, p <0.001, η² = 0.03. A highly significant main effect of trial type was also observed, F(2, 369) = 94.31, p < 0.0001, η² = 0.49, demonstrating that the type of trial accounted for a substantial proportion of the variance in 1/f slope modulation. This large effect suggests that the cognitive or perceptual demands associated with different trial types substantially modulated the 1/f slope. Importantly, the interaction between group and trial type was not statistically significant, F(4, 369) = 0.64, p = 0.65, η² = 0.007. This non-significant interaction indicates that the pattern of trial effects on 1/f slope modulation was consistent across the different groups.

#### 3.3.1. *Post-hoc* analysis: Between trial comparison

As shown in Figure 5, when comparing the modulation of the 1/f slope between standard and non-standard stimuli, we observed a significantly larger change in response to non-standard stimuli (p < 0.001), supporting the hypothesis that greater attention demands elicit stronger modulations in the 1/f slope.

**Figure 5.**
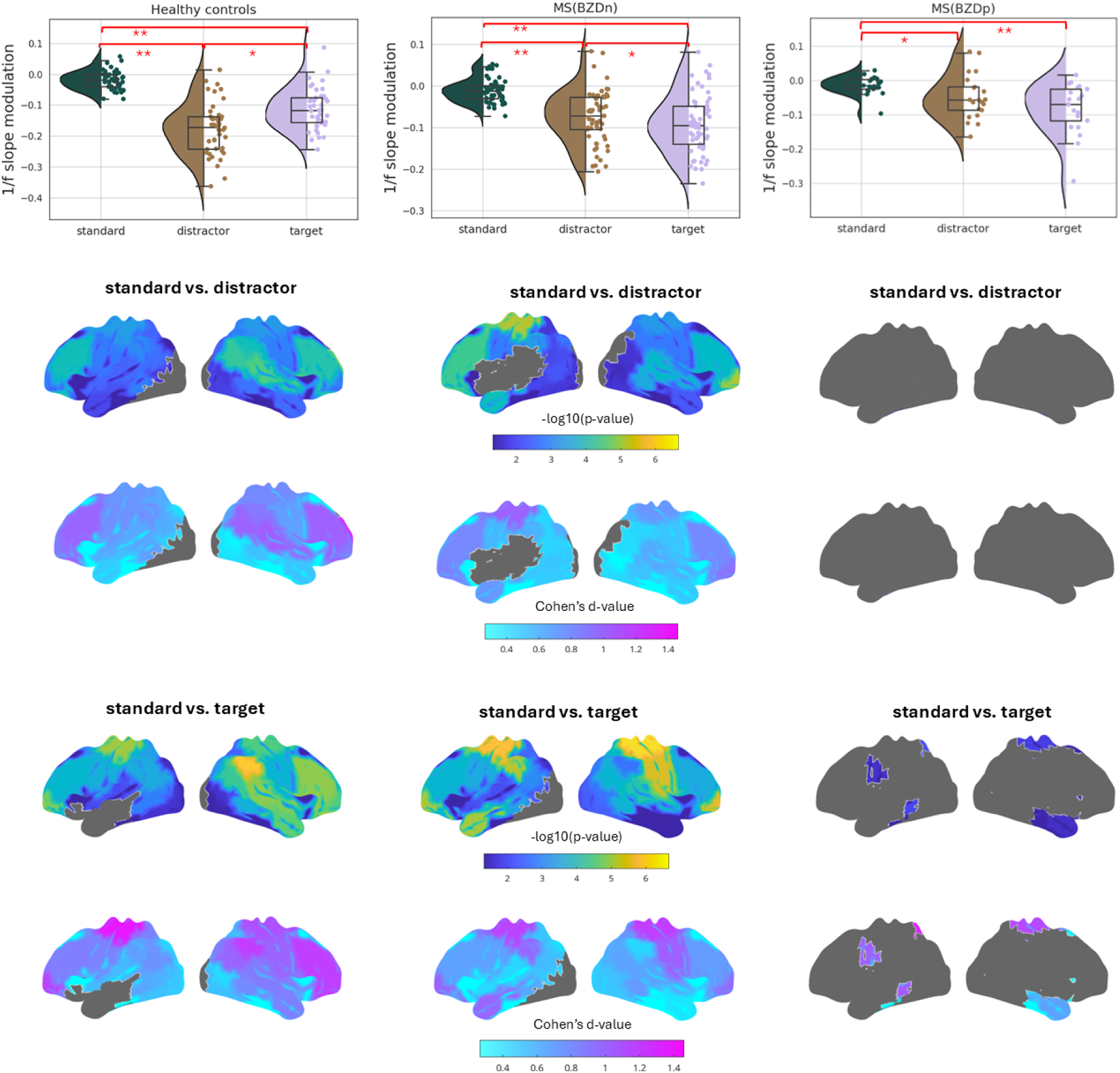
The distribution of average over whole brain 1/f slope modulation for all trials and all groups. The Wilcoxon signed-rank test was used for between-trial comparisons. The spatial distributions of cortical regions show the comparison of 1/f slope modulation between trials at the parcel level (42 brain parcels) after FDR correction for multiple comparisons (**p<0.001, *p<0.05). The colour bars show the value of -10*log(p-value) with significant values higher than 1.3. The spatial distribution of effect size is represented as Cohen’s d-value below each parcel-wise comparison.

In healthy controls, the increase was significantly greater for standard vs distractor (W = 13, Z = -3.57, p < 0.001, d = 0.53) and standard vs target (W = 9, Z = -5.63, p < 0.001, d = 0.84). A similar pattern was observed in pwMS(BZDn), with significant differences between standard vs distractor (W = 100, Z = -6.07, p < 0.001, d = 0.77) and standard vs target (W =68, Z = -6.30, p < 0.001, d = 0.80). In pwMS(BZDp), we similarly found a significant difference between standard vs distractor (W = 56, Z = 1.40, p = 0.03, d = 0.30) and standard vs target (W = 9, Z = -3.70, p < 0.001, d = 0.80).

When comparing standard trials with target and distractor trials, parcel-wise analysis showed that 1/f slope modulation in the sensorimotor cortex was slightly higher during target trials than during distractor trials, potentially reflecting additional motor processes required by the button press response.

Interestingly, HCs showed a larger 1/f slope modulation in response to distractor trials compared to target trials (W = 270, Z = –5.56, p = 0.008, d = 0.83). This effect reversed direction in pwMS(BZDn) (W = 596, Z = –2.51, p = 0.01, d = 0.32), and no effect was observed in pwMS(BZDp) (W = 59, Z = –1.96, p = 0.05, d = 0.42). The parcel-wise comparisons between target and distractor stimuli did not reveal significant differences in any of the groups.

#### 3.3.2. *Post-hoc* analysis: Between group comparison

Figure 6 shows the distribution of 1/f slope modulation averaged over the whole brain across groups separately for different trial types.

**Figure 6.**
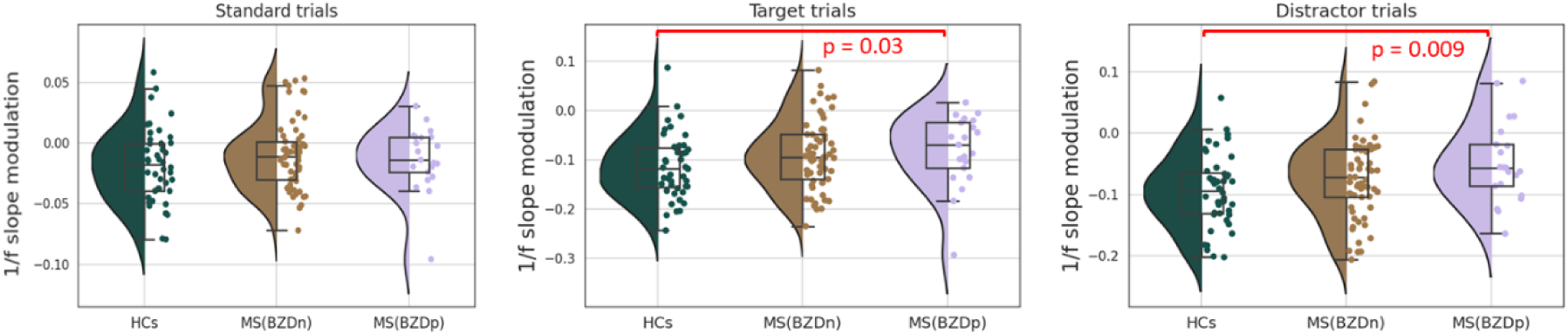
The distribution of 1/f slope modulation averaged over the whole brain across groups separately for different trial types. The Mann-Whitney U test was used for the between-group comparisons.

During standard trials, comparisons between healthy controls (HCs) and pwMS groups did not reach statistical significance (HCs vs. pwMS(BZDn): U = 1152, Z = –1.23, p = 0.21, d = 0.24; HCs vs. pwMS(BZDp): U = 384, Z = –1.08, p = 0.27, d = 0.27). Additionally, no significant differences were observed between the two MS groups (U = 608, Z = –0.33, p = 0.73, d = 0.07).

In contrast, during target and distractor trials, the HCs showed significantly greater modulation compared to pwMS(BZDp): target trials (U = 308, Z = –2.15, p = 0.03, d = 0.55) and distractor trials (U = 277, Z = –2.58, p = 0.009, d = 0.68). However, comparisons between HCs and pwMS(BZDn) were not significant in either target (U = 1092, Z = –1.62, p = 0.10, d = 0.32) or distractor trials (U = 1092, Z = –1.62, p = 0.10, d = 0.32). Additionally, no significant differences were observed between the two MS groups in either target (U = 551, Z = –0.94, p = 0.34, d = 0.21) or distractor trials (U = 512, Z = –1.35, p = 0.17, d = 0.30).

Overall, these results partially support the hypothesis that healthy subjects demonstrate a more pronounced inhibitory response compared to pwMS, even though effects did not survive parcel-wise correction.

### 3.4. The correlation between the 1/f slope modulation and cognitive scores

For pwMS not receiving benzodiazepines, 1/f slope modulation was significantly correlated with BVMT-R scores in both standard (r = -0.31, p = 0.01) and distractor trials (r = -0.37, p = 0.003), and also showed a negative correlation with VGLT scores in standard trials (r = -0.27, p = 0.03). In pwMS receiving benzodiazepines, a significant correlation was found between 1/f slope modulation and BVMT-R scores in distractor trials (r = -0.53, p = 0.02). No significant correlations were observed in healthy controls. Detailed results are provided in Table S1, and corresponding scatter plots are shown in Figure S2.

### 3.5. The correlation between the 1/f slope modulation following an auditory and n-back stimuli

To evaluate whether 1/f slope modulation reflects a consistent, paradigm-independent mechanism, we conducted additional analyses on 101 participants who performed both auditory oddball and the n-back working memory tasks (HCs = 34, pwMS(BZDn) = 50, pwMS(BZDp) = 17). We focused on target and distractor trials in the oddball task to align with the two-condition design (target vs distractor) in the n-back data. We corrected the correlation analyses for the multiple comparisons.

Healthy controls and pwMS(BZDn) showed a significant positive correlation in 1/f slope modulation between the two tasks across all n-back load levels (0-back, 1-back, 2-back), as shown in Figure 7 and Figure 8, respectively. Individuals who showed larger (or smaller) 1/f slope changes in the oddball task tended to demonstrate a similar pattern in the n-back task. Among pwMS(BZDp), correlations did not reach significance for the 0-back and 1-back target and 0-back distractor conditions but exhibited a similar trend, see Figure S3.

**Figure 7.**
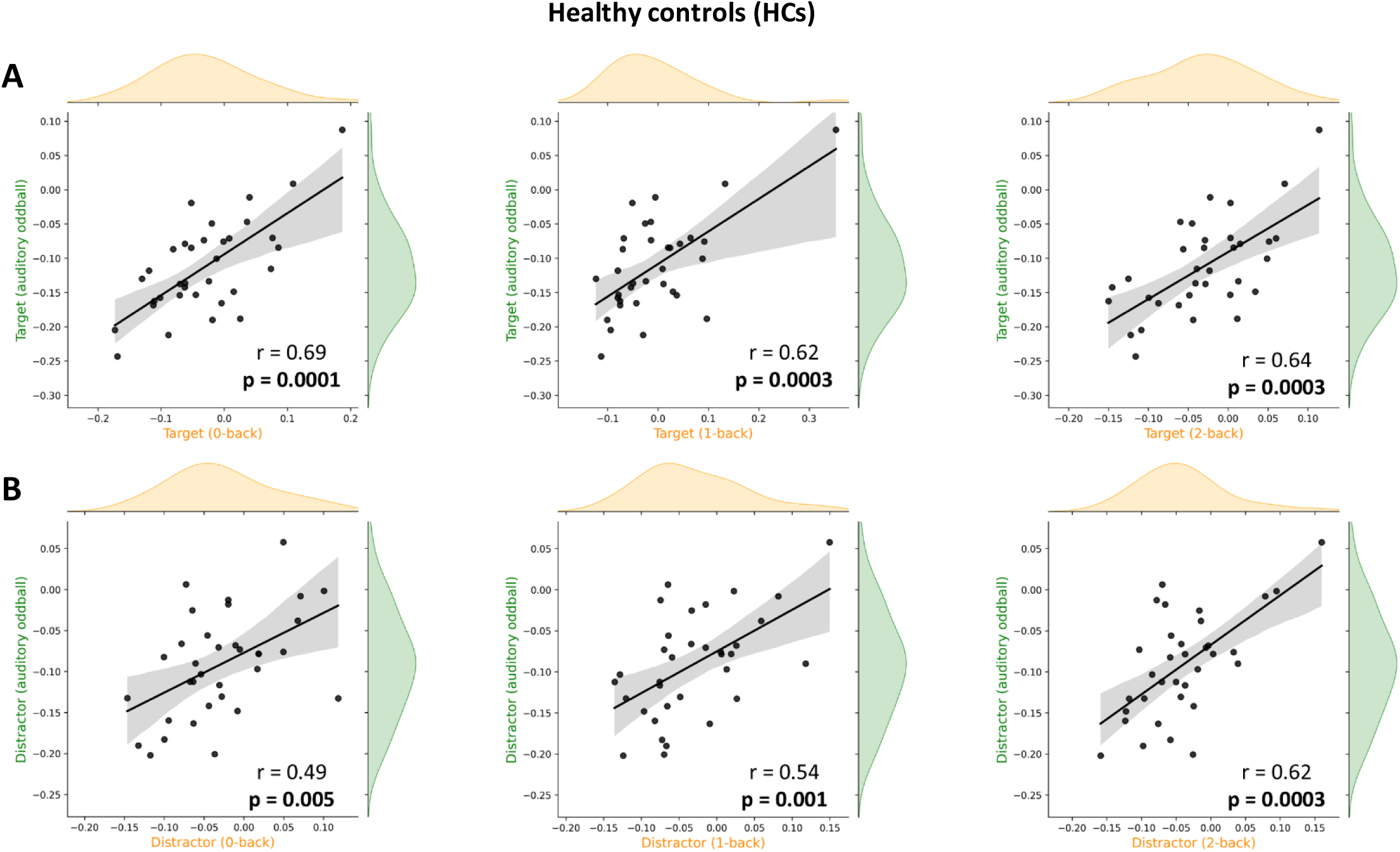
Association of 1/f slope modulation between the auditory oddball and n-back tasks (0-back, 1-back, and 2-back conditions) in **healthy controls** (HCs). Panels **A** and **B** show the results for target and distractor trials, respectively.

**Figure 8.**
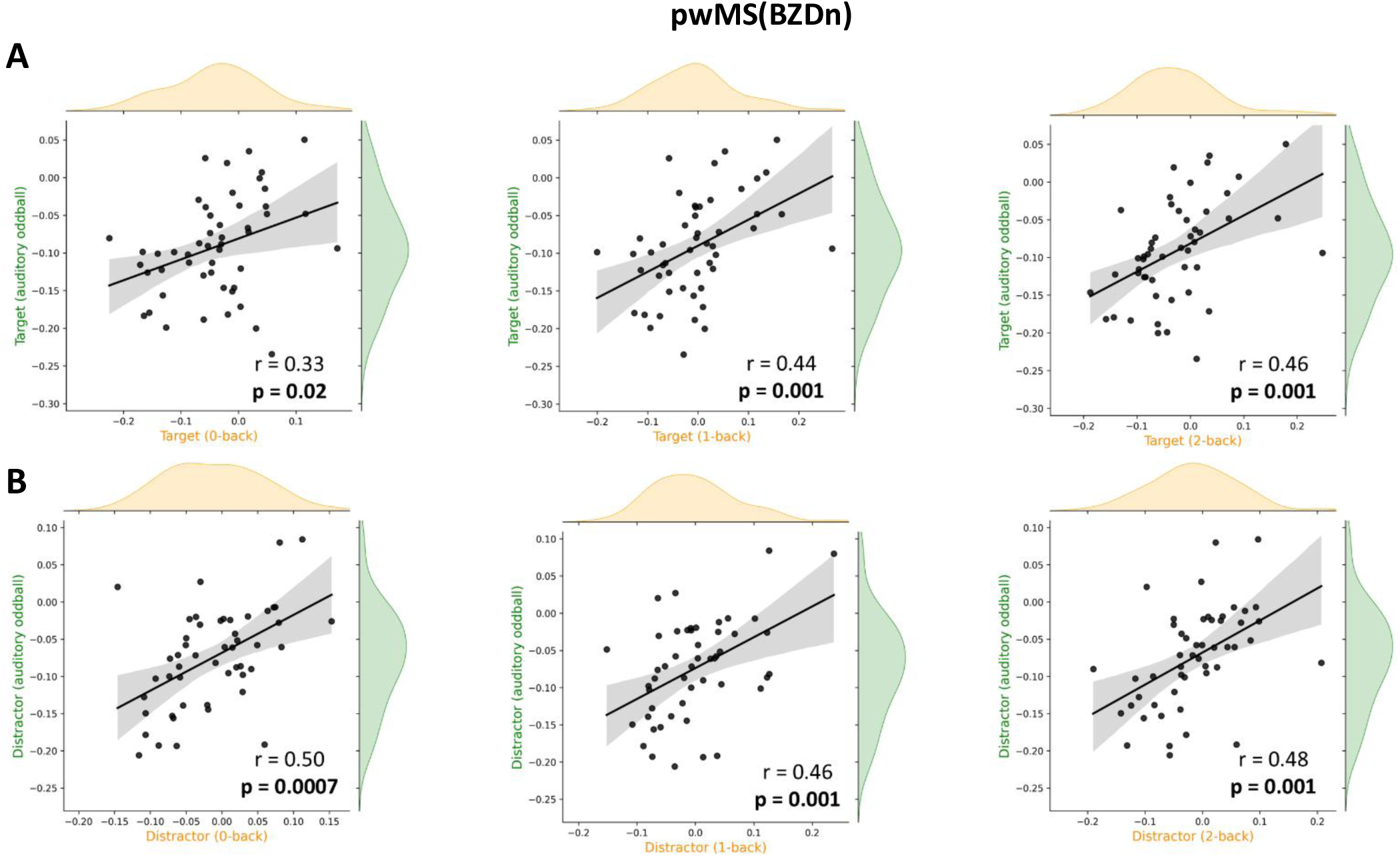
Association of 1/f slope modulation between the auditory oddball and n-back tasks (0-back, 1-back, and 2-back conditions) in **pwMS(BZDn)**. Panels **A** and **B** show the results for target and distractor trials, respectively.

## 4. Discussion

This study explores the neurophysiological underpinnings of cognitive impairments in multiple sclerosis through the lens of the aperiodic 1/f slope modulation by leveraging MEG data during auditory oddball and visual-verbal n-back tasks. We investigated the modulation of the 1/f slope as an indicator of excitation/inhibition (E/I) balance and its potential role as a paradigm-independent marker of cognitive control. Our findings offer several key insights into the neural mechanisms of cognitive dysfunction in MS and underscore the potential utility of aperiodic spectral components in cognitive neuroscience research.

Consistent with our first hypothesis and prior findings (Akbarian et al., 2024; Gyurkovics et al., 2022; Kałamała et al., 2023), we observed a robust post-stimulus steepening of the 1/f slope across all groups. This modulation was more pronounced and spatially widespread for non-standard stimuli (target and distractor) compared to standard stimuli. This finding aligns with existing literature which indicates that non-standard stimuli, demanding higher attentional and cognitive resources, show higher neural engagement (Gyurkovics et al., 2022; Polich, 2007). Significant modulation was observed in the temporal cortex, a key region for auditory processing (Howard et al., 2000; Nourski, 2017), which exhibits increased activity during auditory task engagement. These results substantiate the role of the 1/f slope as a sensitive marker of cognitive load and attentional mechanisms (Lu et al., 2024), extending beyond traditional oscillatory analyses (Buzsáki & Watson, 2012; Cavanagh & Frank, 2014).

Similar to our previous findings (Akbarian et al., 2024), people with MS treated with benzodiazepines showed reduced 1/f slope modulation in response to distractor stimuli compared to HCs. This suggests that benzodiazepine use may disrupt inhibitory neural mechanism (Nicholson et al., 2018), potentially contributing to the cognitive deficits reported in the literature on benzodiazepines (Barker et al., 2004; Zetsen et al., 2022). Also, HCs showed a larger 1/f slope modulation (suggesting more inhibition) following distractor trials compared to target trials while this pattern was reversed in pwMS. The overall observed trend was consistent with a compromised inhibitory response in pwMS, suggesting an underlying imbalance between excitation and inhibition. This finding resonates with prior evidence of synaptic loss and dysregulation of inhibitory control in MS (Huiskamp et al., 2022; Zoupi et al., 2021) and highlights the potential of aperiodic spectral slope in capturing subtle neurophysiological alterations that may underlie cognitive deficits. Importantly, directional changes in the 1/f spectral slope (steepening or flattening) offer a direct and interpretable marker of shifts in E/I balance, in contrast to the often challenging interpretation of ERFs polarity (Hajizadeh et al., 2019).

Also, in line with our previous work (Akbarian et al., 2024), we observed that the task-induced modulation of the 1/f slope exhibited a significant negative correlation with offline visuospatial memory, as measured by the BVMT-R, in pwMS(BZDn). Specifically, a stronger modulation toward inhibition was associated with better visuospatial memory performance, suggesting that enhanced inhibitory neural processes may support cognitive function in this domain. Recent multimodal neuroimaging studies have reported positive correlations between hippocampal GABA concentrations, receptor density, and both visuospatial and verbal memory in pwMS (Huiskamp et al., 2023; Zhang et al., 2024), further emphasizing the role of inhibitory neurotransmission in cognitive performance. Moreover, visuospatial memory has emerged as a critical cognitive domain for distinguishing various cognitive profiles among pwMS (Van Dam et al., 2024), underscoring its clinical relevance. Future research should explore this relationship in greater detail to elucidate the underlying mechanisms, thereby advancing the potential of 1/f slope modulation as a clinical biomarker for cognitive impairment in pwMS.

Several factors may explain the lack of association between 1/f slope modulation and neuropsychological test performance across other groups and cognitive tests. First, the auditory oddball task may impose a relatively modest cognitive load, limiting its sensitivity to detect individual differences in higher-order cognitive functions. Also, it is possible that the 1/f slope modulation captures a distinct aspect of neural function that is not directly reflected in the selected neuropsychological measures. Finally, it is possible that our group-level sample sizes were insufficient to detect a reliable relationship between cognitive performance and slope modulation, particularly given that slope modulation was calculated as a difference score (post-minus pre-exponent), and such difference measures are known to have weaker psychometric properties compared to absolute measures (Hedge et al., 2018; Von Bastian et al., 2020). Future research employing larger samples and a wider array of cognitive tasks varying in complexity may offer a more comprehensive assessment of these relationships.

A salient finding of our study is the significant positive correlation in 1/f slope modulation between the auditory oddball and n-back tasks among HCs and pwMS(BZDn). Subjects who show larger (or smaller) 1/f slope changes in one task tend to show a similar pattern in the other. This cross-task consistency supports that the 1/f slope functions as a trait-like indicator of cognitive control, reflecting underlying E/I balance or top-down regulatory mechanisms engaged across diverse cognitive demands. Recent work using different approaches (multivariate pattern analysis) across three tasks has similarly demonstrated cross-task generalization of 1/f aperiodic components (Lu et al., 2024), further bolstering its role as a paradigm-independent neurophysiological marker. In contrast to the overall pattern observed, the subgroup of pwMS receiving benzodiazepine treatment (pwMS(BZDp)) did not show significant cross-task correlations. The lack of significant findings might be due to the relatively small sample size in the pwMS(BZDp) subgroup, which might have limited the statistical power needed to detect subtle effects.

While our study provides novel insights, some limitations must be acknowledged. One potential limitation of our study is the restricted time window around stimulus onset. Due to the constraints of our task paradigm, we confined the analysis to a 500 -millisecond window, which reduced the frequency resolution to 2 Hz. Despite this, power spectra were modelled with high model fit during aperiodic parametrization in all our analyses (See supplementary materials, Figure S4). Future studies may benefit from extending the time window to improve frequency resolution, with the caveat that this would lead to decreased temporal resolution which may pose a problem if aperiodic changes are relatively short-lived. It is also important to note that while the 1/f slope is often used as an indirect measure of the excitatory/inhibitory (E/I) balance, alternative explanations have been suggested. For example, one theory argues that 1/f scaling may result from fluctuations in the transition rates between cortical up and down states (Baranauskas et al., 2012), whereas another attributes it to the damping effects observed in harmonic oscillators (Muthukumaraswamy & Liley, 2018).

Building on our findings, future research should explore several avenues to deepen our understanding of E/I balance disruptions in pwMS. Longitudinal studies which track changes in 1/f slope modulation over time could elucidate the progression of these disruptions and their relationship with cognitive decline. In addition, employing a broader spectrum of cognitive paradigms with varying demands would better capture the relationship between 1/f slope modulation and specific cognitive domains. Integrating aperiodic spectral measures with structural and functional imaging biomarkers may enhance the predictive power for cognitive impairments in MS. Moreover, employing a variety of preprocessing and analytical methods can help assess the robustness of the observed effects across different methodological choices, such as source reconstruction techniques, that may impact the shape of the power spectrum.

## 5. Conclusion

This study underscores the potential of the aperiodic 1/f slope as a sensitive and paradigm-independent neurophysiological marker of cognitive control and E/I balance. Attenuated distractor-related modulation among people with MS taking benzodiazepines (pwMS(BZDp)) suggests that benzodiazepine use may impair inhibitory neural mechanisms underlying cognitive deficits. The observed correlations across distinct cognitive tasks suggest the utility of the 1/f slope in capturing stable aspects of cognitive processing. These findings pave the way for future research using aperiodic spectral components to better understand cognitive impairments in MS and other neurological conditions.

## Data and Code Availability

Data and code are available at reasonable request from the corresponding author.

## Author contributions

Conceptualization (F.A., M.G., M.B.D., M.D., G.N., J.V.S); Methodology (F.A., M.G., G.N., J.V.S); Software (F.A.); Formal Analysis (F.A., J.V.S.); Original Draft (F.A.); Review & Editing (F.A., M.G., M.B.D., M.D., G.N., J.V.S); Visualization (F.A.); Supervision (G.N., J.V.S); and Funding Acquisition (G.N., J.V.S).

## Acknowledgements

The authors thank all participants for their enthusiasm and commitment to participate.

## Supplementary Materials

**Figure S1.**
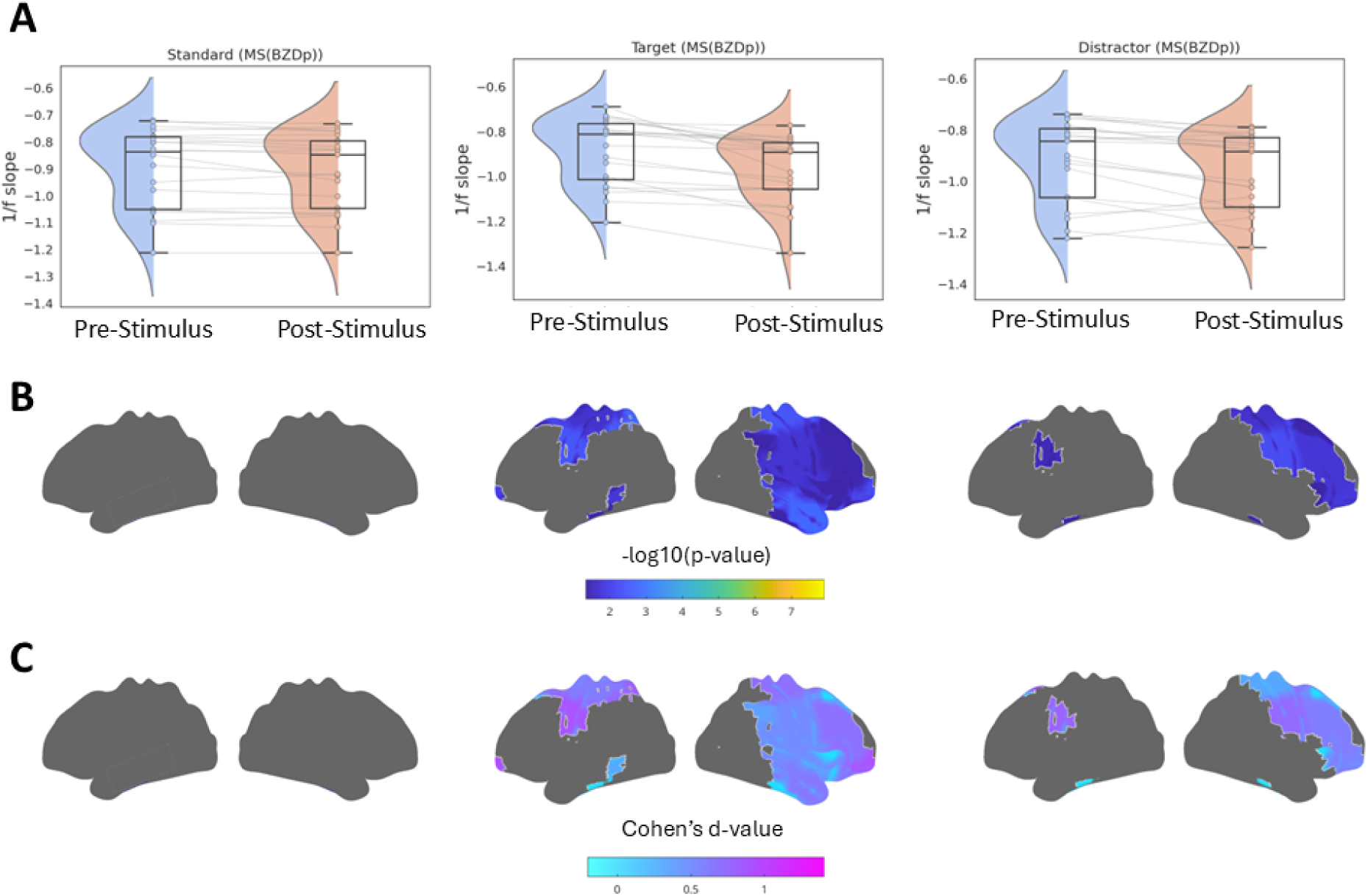
**(A)** Distribution of whole-brain averaged 1/f slope values measured pre- and post-stimulus for all trials in **pwMS(BZDp)**. The Wilcoxon signed-rank test yielded p-values < 0.0001 for all within-group comparisons. **(B)** Spatial distribution maps at the parcel level (42 brain parcels) comparing the 1/f slope pre- and post-stimulus, with corrections for multiple comparisons using the FDR method. **(C)** Spatial distribution of effect sizes, represented as Cohen’s d for each parcel-wise comparison.

**Table S1.** Results of correlation analysis between the 1/f slope modulation and cognitive scores.

|  | Standard trial | Target trials | Distractor trials |
| --- | --- | --- | --- |
| **BVMT-R** | HCs: $r = 0.14$, $p = 0.35$<br>**pwMS(BZDn): $r = -0.31$, $p = 0.01$**<br>pwMS(BZDp): $r = 0.05$, $p = 0.84$ | HCs: $r = 0.08$, $p = 0.58$<br>pwMS(BZDn): $r = -0.23$, $p = 0.07$<br>pwMS(BZDp): $r = -0.27$, $p = 0.27$ | HCs: $r = 0.23$, $p = 0.11$<br>**pwMS(BZDn): $r = -0.37$, $p = 0.003$**<br>**pwMS(BZDp): $r = -0.53$, $p = 0.021$** |
| **SDMT** | HCs: $r = -0.15$, $p = 0.31$<br>pwMS(BZDn): $r = -0.21$, $p = 0.09$<br>pwMS(BZDp): $r = 0.21$, $p = 0.39$ | HCs: $r = 0.001$, $p = 0.99$<br>pwMS(BZDn): $r = -0.04$, $p = 0.72$<br>pwMS(BZDp): $r = 0.08$, $p = 0.74$ | HCs: $r = 0.01$, $p = 0.90$<br>pwMS(BZDn): $r = -0.17$, $p = 0.17$<br>pwMS(BZDp): $r = -0.03$, $p = 0.87$ |
| **VGLT** | HCs: $r = 0.14$, $p = 0.36$<br>**pwMS(BZDn): $r = -0.27$, $p = 0.03$**<br>pwMS(BZDp): $r = -0.03$, $p = 0.87$ | HCs: $r = 0.08$, $p = 0.56$<br>pwMS(BZDn): $r = -0.16$, $p = 0.19$<br>pwMS(BZDp): $r = -0.24$, $p = 0.33$ | HCs: $r = 0.19$, $p = 0.19$<br>pwMS(BZDn): $r = -0.25$, $p = 0.05$<br>pwMS(BZDp): $r = -0.09$, $p = 0.70$ |
| **COWAT** | HCs: $r = -0.20$, $p = 0.19$<br>pwMS(BZDn): $r = -0.08$, $p = 0.52$<br>pwMS(BZDp): $r = -0.19$, $p = 0.43$ | HCs: $r = 0.005$, $p = 0.97$<br>pwMS(BZDn): $r = -0.12$, $p = 0.33$<br>pwMS(BZDp): $r = -0.23$, $p = 0.34$ | HCs: $r = -0.01$, $p = 0.91$<br>pwMS(BZDn): $r = -0.15$, $p = 0.22$<br>pwMS(BZDp): $r = -0.28$, $p = 0.25$ |

**Figure S2.**
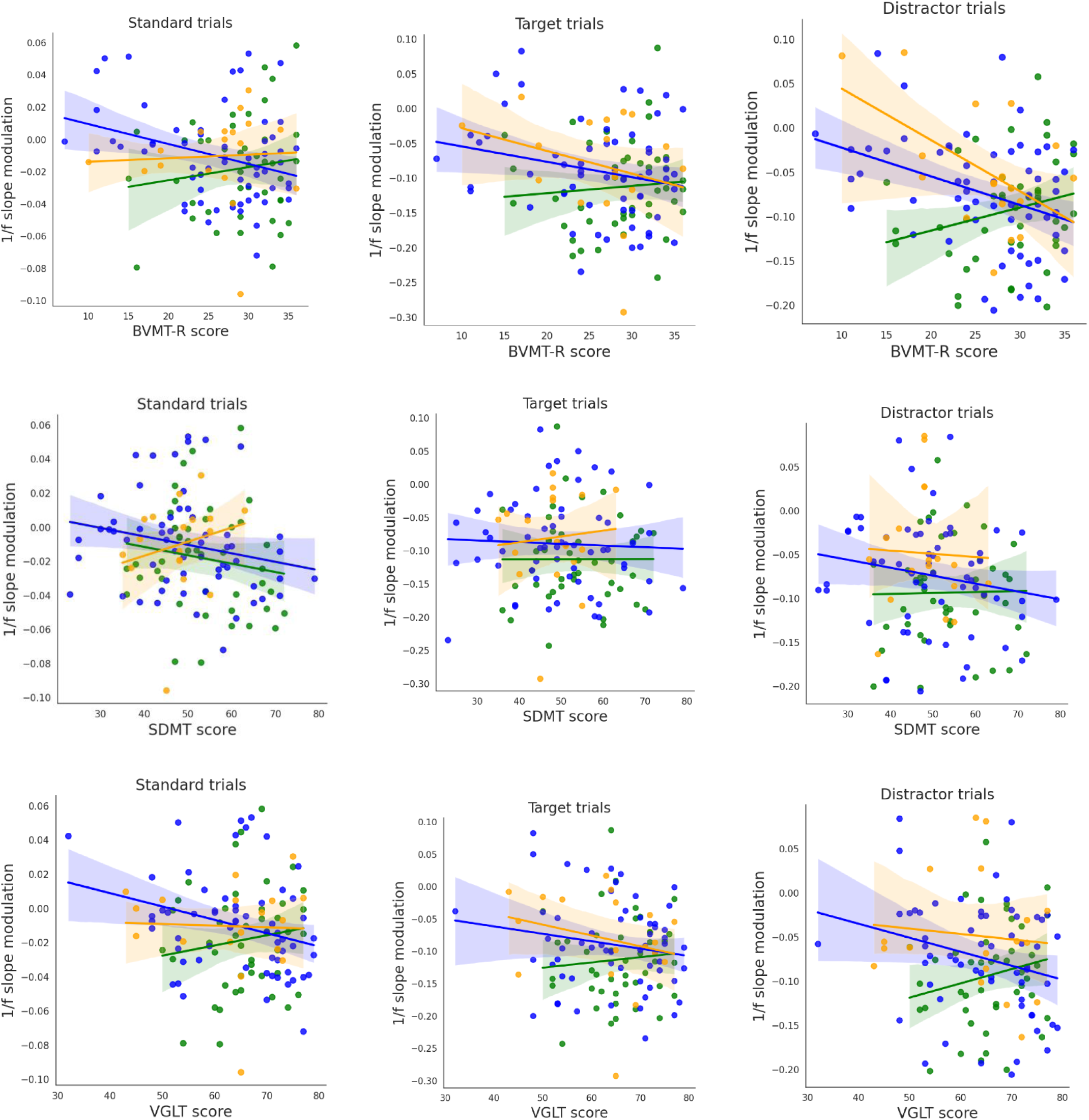
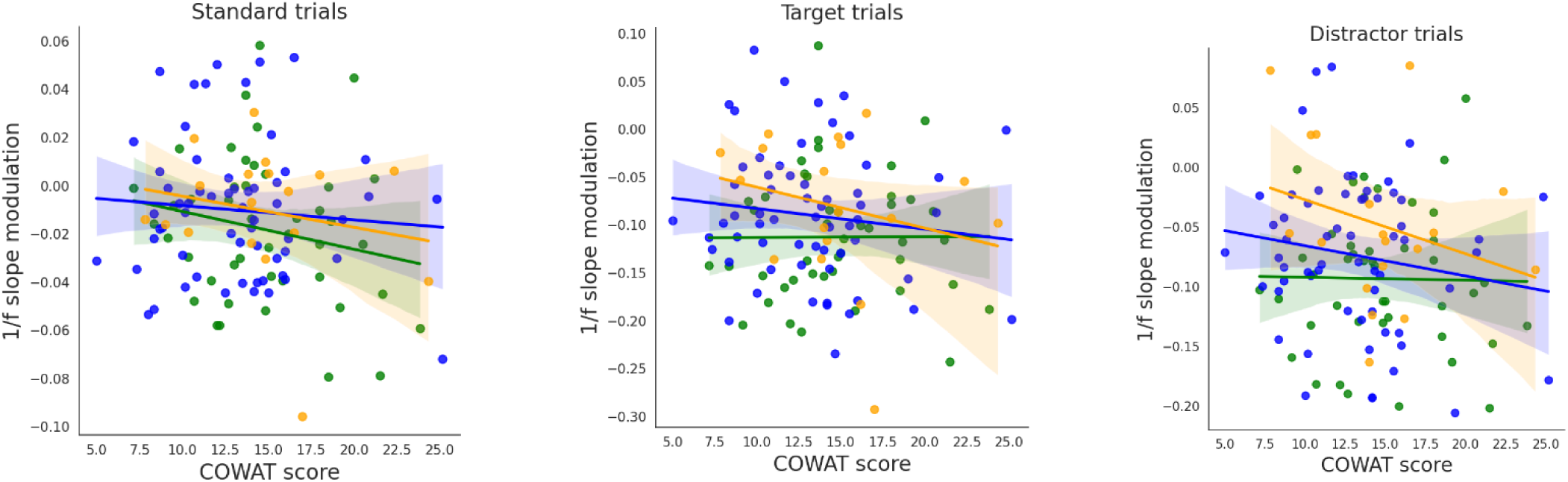
Scatter plots of correlation analysis between the 1/f slope modulation and cognitive scores for all subjects. Green, blue and orange dots represent healthy controls, pwMS(BZDn) and pwMS(BZDp), respectively. BVMT-R (Revised Brief Visuospatial Memory Test: to assess visuospatial memory. SDMT (Symbol Digit Modalities Test): to evaluate information processing speed. VGLT (the Dutch version of the California Verbal Learning Test (CVLT-II)): to assess verbal memory. COWAT (Controlled Oral Word Association Test): to measure verbal fluency.

**Figure S3.**
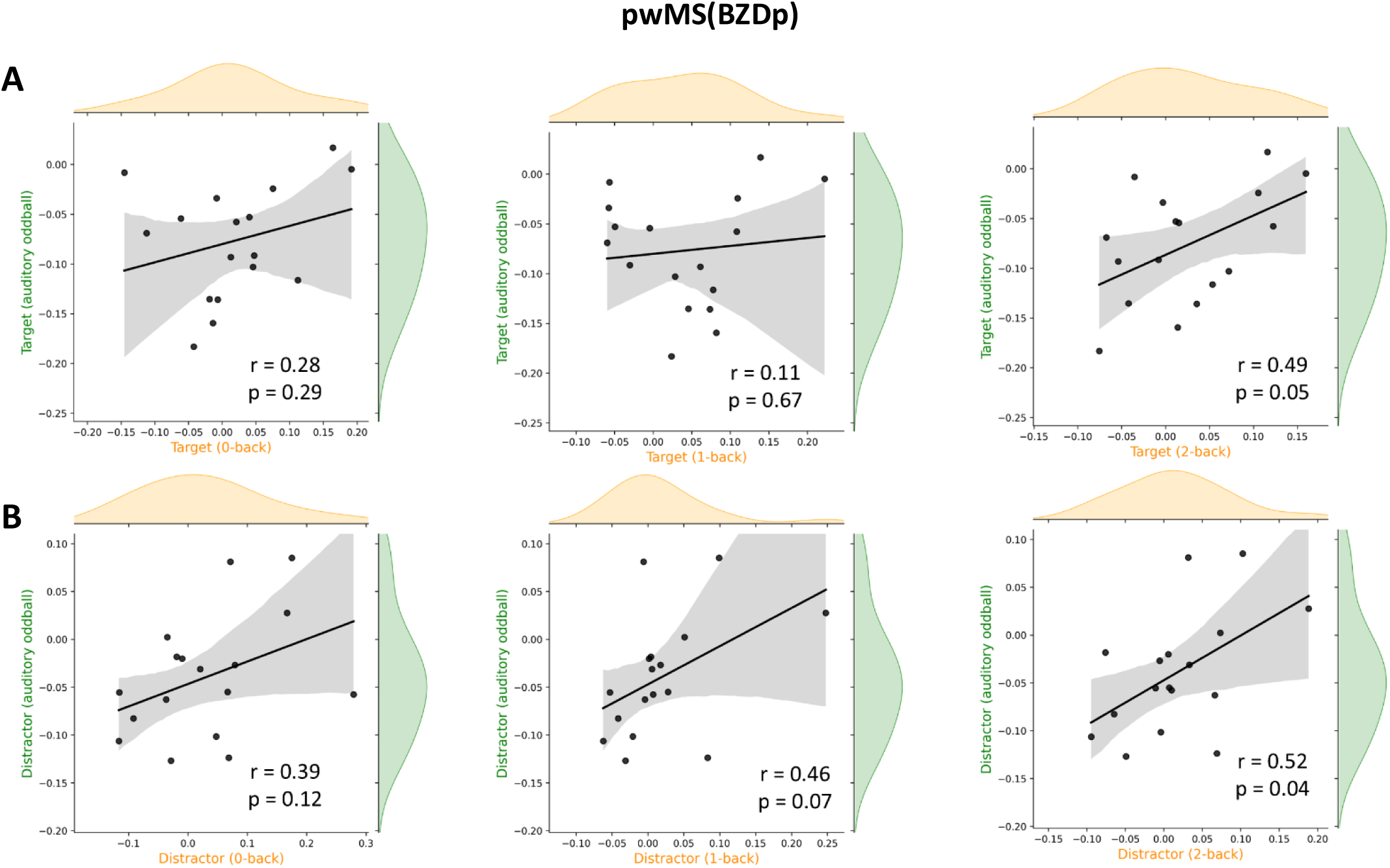
Association of 1/f slope modulation between the auditory oddball and n-back tasks (0-back, 1-back, and 2-back conditions) in **pwMS(BZDp)**. Panels **A** and **B** show the results for target trials and distractor trials, respectively.

**Figure S4.**
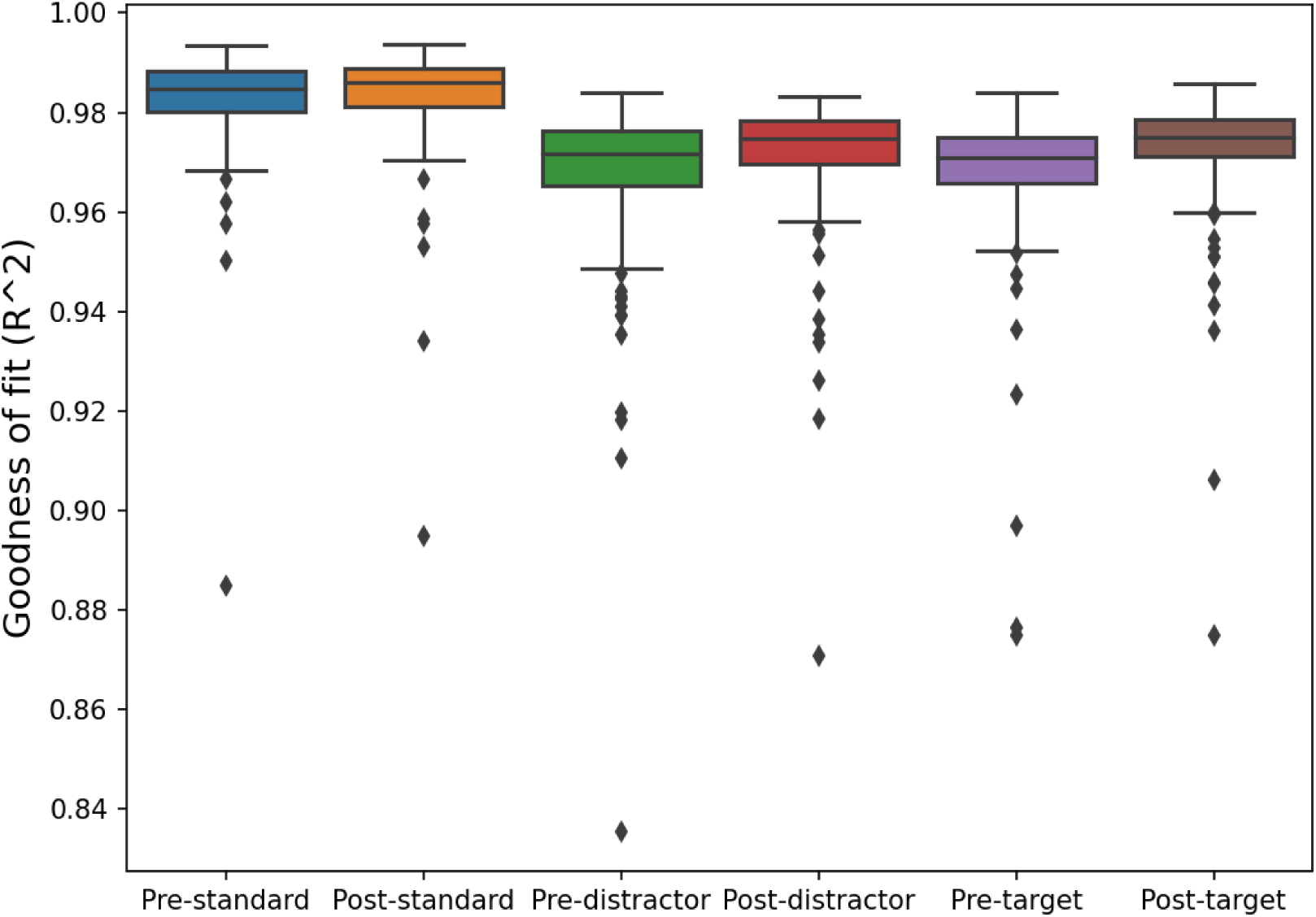
Goodness of Fit. Box plots of the R2 parameter of FOOOF fitting of power spectrum densities within pre- and post-stimulus time windows for each type of trial for all subjects.

